# Persistence of *Plasmodium cynomolgi* hypnozoites in cynomolgus monkey iPS-derived hepatocytes

**DOI:** 10.1101/2021.11.16.468833

**Authors:** Mélanie Pellisson, Anne-Marie Zeeman, Thierry Doll, Lucy Kirchhofer-Allan, Sven Schuierer, Guglielmo Roma, Erika L. Flannery, Sebastian A. Mikolajczak, Clemens H. M. Kocken, Pascal Maeser, Matthias Rottmann, Matthias Mueller

## Abstract

*Plasmodium cynomolgi (Pc)* is one of the few parasite species that forms quiescent liver stage parasites known as hypnozoites and is therefore a suitable model for *Plasmodium vivax*. Very little is known about liver stage dormancy, which hampers the search for compounds with anti-hypnozoite activity. Here, we present the development of a *Pc in vitro* infection model using stem cell-derived hepatocytes from *Macaca fascicularis*. IPS cells were established on feeder free condition and differentiated into hepatocytes via inducible overexpression of key transcription factors. The generated hepatocytes were infected with *Pc* sporozoites and hypnozoite formation as well as schizont development were confirmed by immunofluorescence. This system is a promising tool to study the mechanisms underlying liver stage dormancy and facilitate drug discovery against hypnozoites.

## Main Text

In 2018, malaria had an estimated global incidence of 228,000,000 and a death toll of 405,000 (WHO, 2019). Thus, in spite of the recent successes in the fight against malaria (Rabinovich et al., 2017), the disease remains to be a main killer in the tropics. Malaria being anthroponotic, elimination and even eradication are theoretically possible. However, in addition to drug-resistant *Plasmodium* parasites and insecticide-resistant *Anopheles* mosquitoes (Ashley, Pyae Phyo, & Woodrow, 2018), there is an ultimate hindrance to malaria eradication: the hypnozoites (Campo, Vandal, Wesche, & Burrows, 2015). Hypnozoites are small, uninucleated, intracellular stages that stay dormant for months or even years in the liver of infected carriers (Dembele et al., 2014; Krotoski et al., 1982), building a hidden reservoir of parasites that, upon reactivation, causes relapsing malaria. Primaquine and tafenoquine are the only drugs that eliminate hypnozoites (Llanos-Cuentas et al., 2019), but their use is limited due to severe side effects in glucose-6-phosphate dehydrogenase-deficient patients. Therefore, new molecules with anti-hypnozoite activity are needed.

Hypnozoite biology is elusive and the mechanisms underlying dormancy and reactivation remain to be elucidated. Most studies on *Plasmodium vivax* (*Pv)* hypnozoites have been conducted with human primary hepatocytes as host cells. These, however, present problems regarding quality, donor-to-donor variability, and ethical considerations. Moreover, working with *Pv* is challenging as there is no long term *in vitro* culture system and sporozoite sourcing is extremely limited (Tachibana et al., 2012). The best surrogate organism for *Pv* is *Plasmodium cynomolgi (Pc)*, a closely related and more accessible malaria parasite of non-human primates that also produces hypnozoites (Krotoski et al., 1980). Here we present for the first time a solution that overcomes the limitations of both *Pv in vitro* work and primary host cell usage by establishing an *in vitro* system for *Pc* hypnozoites that uses cynomolgus monkey hepatocyte-like cells generated by iPS (induced pluripotent stem cells) technology.

Cynomolgus iPS (cynIPS) cells were generated from dermal fibroblasts from two different monkeys by reprogramming, using Sendai viruses bearing the transcription factors Oct3/4, Sox2, Klf4, and c-Myc (Takahashi & Yamanaka, 2006). Pluripotent colonies appeared two to three weeks after infection and were kept on irradiated mouse embryonic fibroblasts. After several passages, the iPS cells were transferred on Matrigel, but spontaneous differentiation occurred on this feeder-less system (Figure 1-figure supplement 1A). Further investigations showed that a higher concentration of bFGF (basic fibroblast growth factor) was essential to maintain pluripotency (Figure 1-figure supplement 1B). Following the clonal expansion phase, clones exhibiting typical human embryonic stem cell-like morphology (flat and tightly packed colonies of round cells with large nuclei) were selected and characterized by immunofluorescence (Figure 1-figure supplements 2 and 3). Based on this result the best clone (#13) was selected and checked for karyotype (Figure 1-figure supplement 4), and differentiation potential into the three germ layers (Figure 1-figure supplement 5 and movie 1).

The generation of human hepatocytes from stem cells is a well-known process and diverse protocols have been developed (Chen et al., 2012; Gao et al., 2017; Si-Tayeb et al., 2010), but the translation of this process to cynomolgus cells required extensive tailoring. The stepwise approach, which consists of adding growth factors mimicking liver embryogenesis, was not robust and reproducible enough (data not shown). Therefore, we used a direct differentiation approach to generate cynomolgus hepatocyte-like cells (cynHLCs) by overexpressing essential transcription factors for hepatic lineage formation (Huang et al., 2014). Using an inducible construct based on a tetracycline ON/OFF system (Figure 1-figure supplement 6), HNF4A (hepatocyte nuclear factor 4 homeobox A) and FOXA3 (forkhead box protein A3) were overexpressed in cynIPS cells to drive endoderm formation and hepatocyte maturation (Kang et al., 2018; Yoon et al., 2018). Briefly, confluent cynIPS cells were induced for 4 days with doxycycline (Dox), and hepatic progenitors were further matured for 2 days to generate cynHLCs (Figure 1A). CynHLCs produced from clone HNF4A/FOXA3-#13.5 displayed the best hepatic characteristics, such as ALB (albumin), HNF1A (hepatocyte nuclear factor 1 homeobox A), HNF4A, A1AT (α1-antitrypsin), KRT18 (cytokeratin 18), AFP (α-fetoprotein), and FOXA2 (Forkhead Box A2) expression (Figure 1B and Figure 1-figure supplement 7) and high levels of albumin secretion (Figure 1C). Additional characterization methods revealed accumulation of lipid droplets (Oil Red O staining) and glycogen storage (Periodic Acid Schiff staining), which are attributes of hepatocytes (Figure 1D).

**Figure 1.**
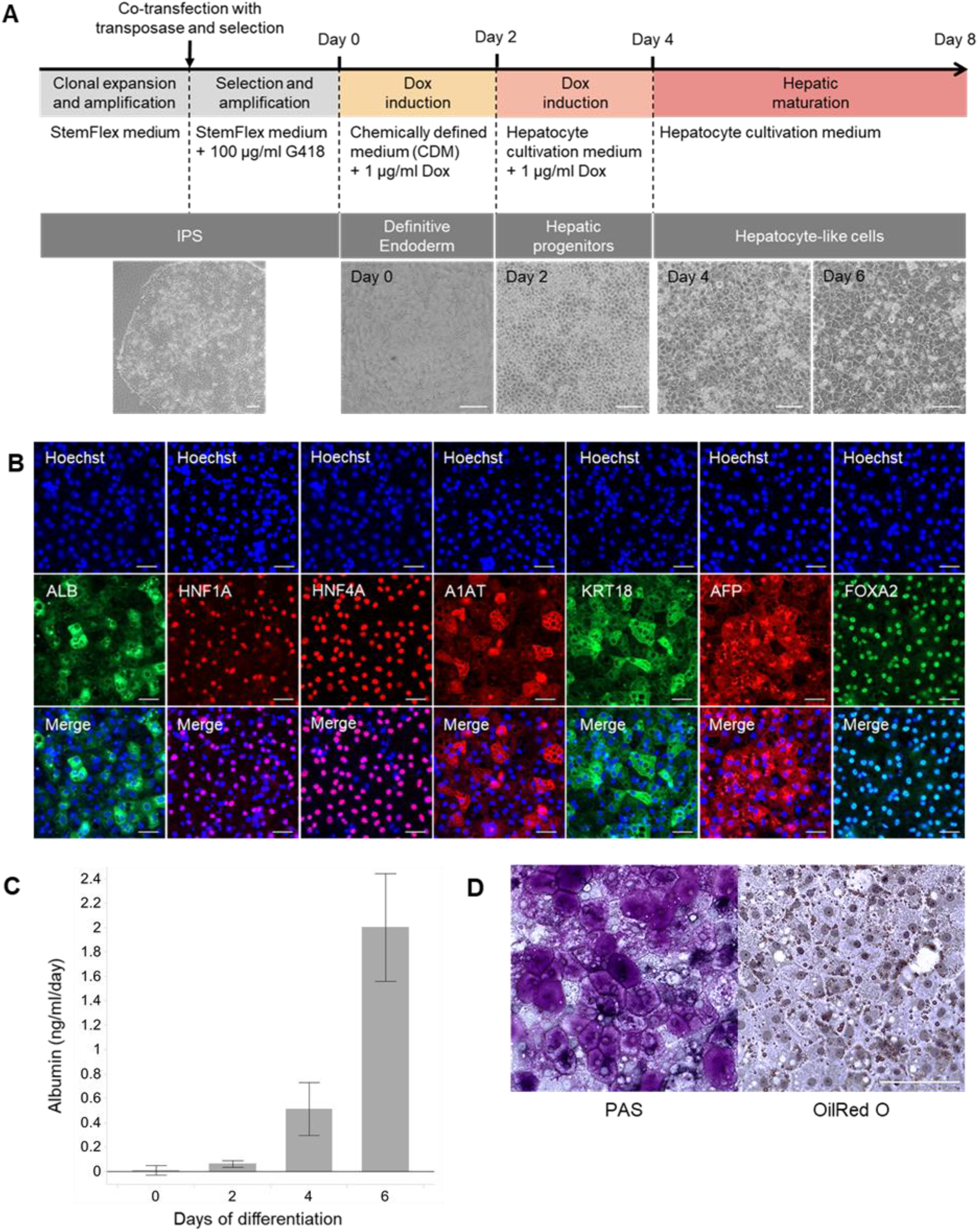
Generation and characterization of cynHLCs. **A**, Schematic representation of the differentiation protocol and brightfield images of differentiating cynIPS cells at the indicated time points. G418: geneticin. Scale bars, 100 µm. **B**, Immunofluorescence of cynHLCs at day 6 with antibodies to HNF1A, HNF4A, AFP, A1AT (red), ALB, KRT18, and FOXA2 (green). Nuclei were visualized with Hoechst (blue). Scale bars, 50 μm. **C**, Enzyme-linked immunosorbent assay (ELISA) for ALB in the supernatants at indicated time points. n=3. **D**, PAS (Periodic Acid Schiff) and Oil Red O staining of cynHLCs after 10 days of differentiation. Scale bars, 100 μm.

RNA-seq analysis of cynHLCs confirmed the hepatic signature of the generated cells (Figure 2A). Comparison with human HLCs as well as liver cancer cell lines (HepG2 and Huh7) showed a high similarity on the mRNA expression profile. While differentiating, the obtained cynHLCs lost expression of pluripotent markers and expressed typical hepatic progenitor genes and mature hepatic genes including efflux transporters at day 7. We also assessed the expression of known *Plasmodium* host entry receptors such as CD81 (Cluster of Differentiation 81) and SRB1 (Scavenger Receptor class B type 1) in the generated cynHLCs. Expression of SRB1 and CD81 was confirmed in cynHLCs at the mRNA level during the differentiation process (Figure 2B). CD81 was gradually expressed in iPS (day 0) and differentiating cells. In contrast, SRB1 was weakly expressed in iPS cells but highly and constantly after day 3, suggesting that cynHLCs become permissive to *Plasmodium* only when they acquire a hepatic signature (Ng et al., 2015). Therefore, cynHLCs were infected when they expressed hallmark hepatic markers like ALB, HNF4A, and AFP in addition to the host entry receptors SRB1 and CD81 (Figure 2C). Although SRB1 was already highly expressed at day 3, sporozoite inoculation was performed at day 4 to avoid potential parasite inhibition due to doxycycline (Gaillard, Madamet, & Pradines, 2015; Pang, Limsomwong, & Singharaj, 1988).

**Figure 2.**
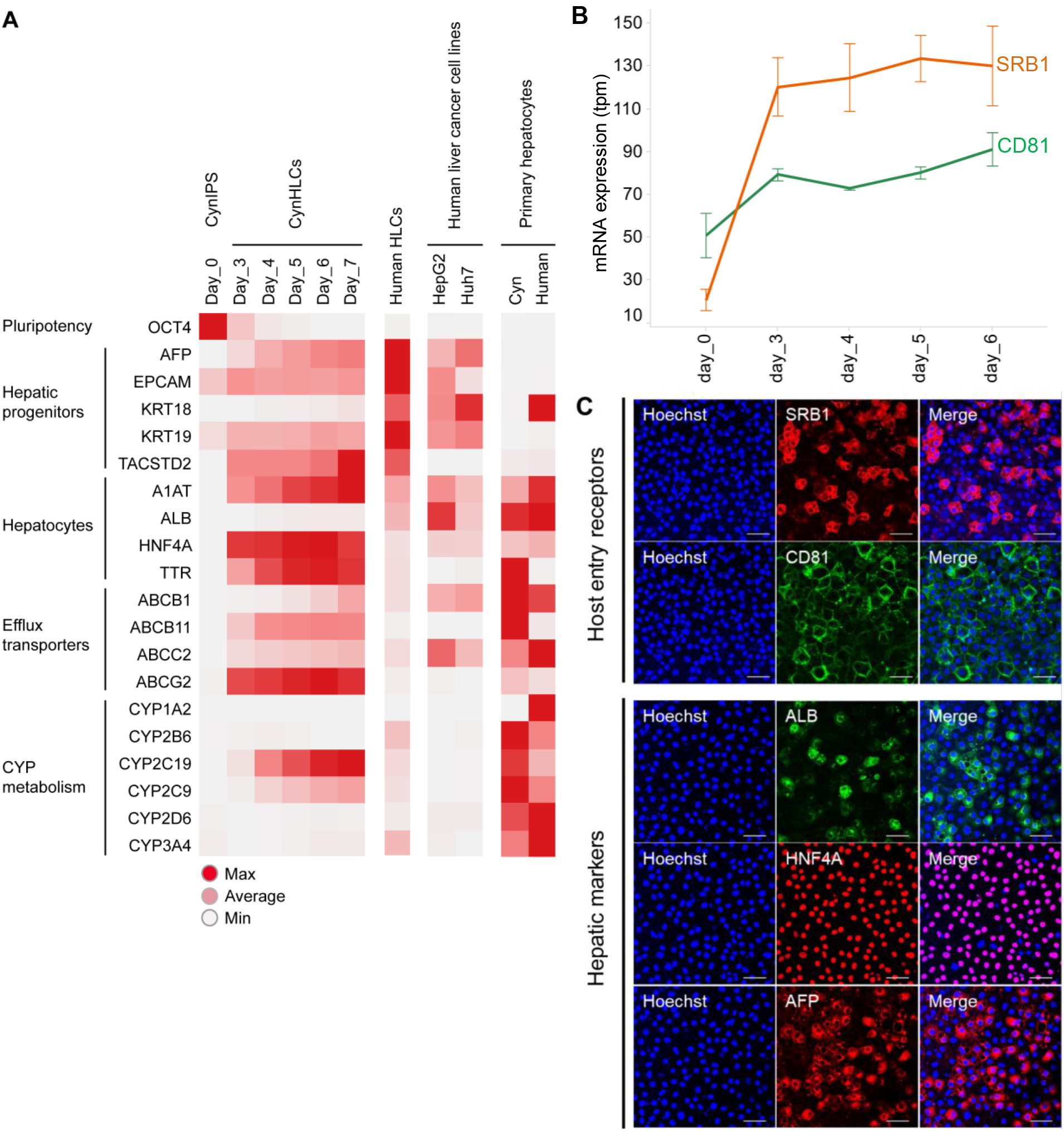
Hepatic signature and expression of host entry receptors indicate permissiveness of cynHLCs to *Plasmodium* parasite. **A**, Heatmap of the RNA-seq based mean expression of cynIPS cells (Day_0), cynHLCs (Day_3 to Day_7), human HLCs, human liver cancer cell lines (HepG2 and Huh7), and primary hepatocytes for selected genes indicative of cell types (Pluripotency, Hepatic progenitors, and Hepatocytes) or cellular processes (Efflux transporters and CYP metabolism). Expression values in transcripts per million (tpm) are colored relative to the maximum (red) and minimum (white) expression value for each gene. **B**, Line chart representing mRNA levels in tpm of SRB1 and CD81 at day 0, 3, 4, 5, and 6. Error bars represent standard deviation. n=3. **C**, Immunofluorescence of cynHLCs at day 4 with antibodies to the hepatic markers ALB (green), HNF4A (red), and AFP (red) and to the host entry receptors SRB1 (red) and CD81 (green). Nuclei were visualized with Hoechst (blue). Scale bars, 50 μm.

While the differentiation process was extremely fast with the cynomolgus cells, the maintenance of the hepatic stage was challenging. After 8 days of differentiation, cynHLCs started to detach and died. In order to improve cell survival, we conducted a screen against an in-house chemical library of well-characterized compounds. Three hit compounds that improved cell viability were identified: a p38 MAPK (mitogen-activated protein kinase) inhibitor named Doramapimod, a DHODH (dihydroorotate dehydrogenase) inhibitor, and a Raf inhibitor (Figure 3-figure supplement 1). Daily treatment with a mix containing the three hits enhanced hepatocyte survival up to 17 days of differentiation (Figure 3A) and maintained ALB and HNF4A expression (Figure 3B). With the triple combination, cell viability remained constant from day 6 to day 12 whereas it dramatically decreased with untreated cells (DMSO control) or cells treated with only one hit compound (Figure 3C). Although the treated cynHLCs showed a more precursor-like mRNA signature than untreated cynHLCs (Figure 3-figure supplement 2), the triple combination allows now to study liver stage (LS) parasite development for at least 12 days, a timeframe necessary to investigate the dormant stage of malaria.

**Figure 3.**
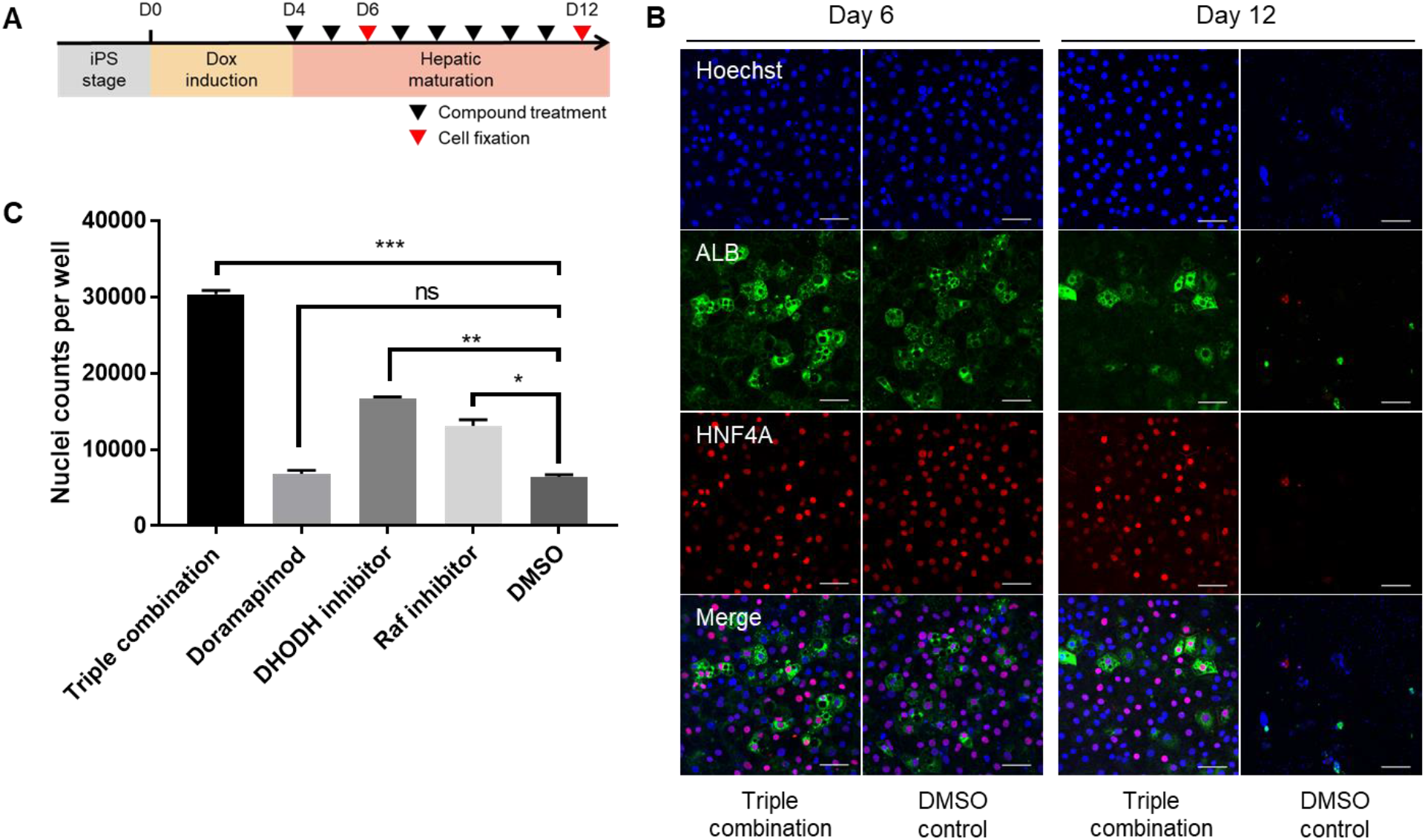
Survival screen allows identification of compounds that improve cynHLCs longevity. **A**, Schematic representation of the experiment conducted to assess the effect of hit compounds identified to promote cynHLCs survival. Cells were treated daily from day 4 (D4) to day 12 (D12). **B**, Immunofluorescence at day 6 and 12 of treated (triple combination) and untreated (DMSO control) cynHLCs with antibodies to ALB (green) and HNF4A (red). Nuclei were visualized with Hoechst (blue). Scale bars, 50 μm. **C**, Bar chart representing the nuclei counts of treated and untreated cynHLCs at day 12. Error bars represent standard deviation. n=3.

Infection of 4-days differentiated cynHLCs was performed with freshly isolated *Pc* sporozoites from *Anopheles stephensi* mosquitoes. We detected the first LS parasites at 2 days post-infection (dpi) after fixation and staining with *Pc* anti-Hsp70 (70 kilodalton heat shock proteins) antibodies. An additional staining with antibodies against UIS4 (up regulated in infective sporozoites gene 4) confirmed that the parasitophorous vacuole membrane (PVM) of LS parasites was formed (Bertschi et al., 2018). At 2 dpi, we observed a uniform population of uninucleated parasites with a size of 3 µm (Figure 4A). Despite the low infection rate of less than 10 parasites per well in a 96 well plate, two populations of LS parasites were clearly distinguishable by 4 dpi. The first (Figure 4B) was composed of stationary parasites that stayed uninucleated over time (Dembele et al., 2014). These parasites were defined as hypnozoites based on their small size (diameter < 8 µm) and their unique nucleus. The second population instead (Figure 4C) was composed of growing and multinucleated forms with a larger diameter, which would lead to the formation of liver schizonts. Developing schizonts and hypnozoites were observed until 12 dpi. As previously described (Mikolajczak et al., 2015), the hypnozoites were slightly increasing in size (from 3 µm at 4 dpi to 5 µm at 12 dpi), but the single nucleus observed by Hoechst staining confirmed the quiescent state of these LS parasites. The current conditions did not allow us to visualize hypnozoite reactivation and subsequent development into mature liver schizonts as described for *Pc* in primary cynomolgus hepatocytes (Dembele et al., 2014).

**Figure 4.**
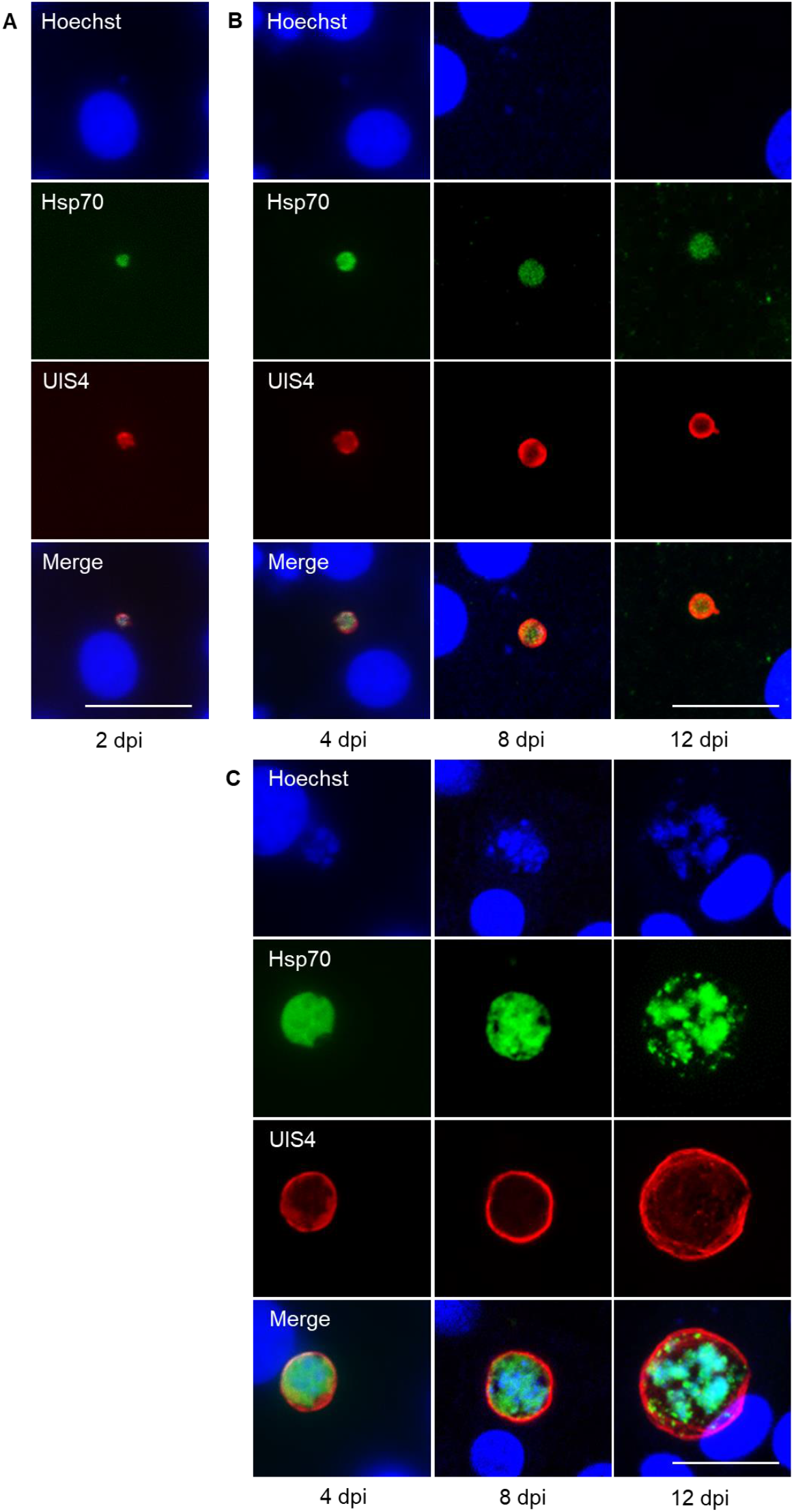
Infection of cynHLCs with *Pc* sporozoites. Immunofluorescence of liver stage parasites with antibodies specific for *Pc*-Hsp70 (green) and *Pc*-UIS4 (red) at 2 dpi (**A**), 4-12 dpi small forms (**B**) and 4-12 dpi large forms (**C**). Cell nuclei and parasite DNA were visualized with Hoechst (blue). Scale bars, 20 µm.

In summary, we report for the first time the generation of cynHLCs from the *Pc* natural host *Macaca fascicularis*. Infection of these host cells with *Pc* sporozoites resulted in successful formation of hypnozoites and persistence until 12 dpi. Moreover, combining the iPS technology with *in vitro* cultured *Pc* erythrocytic stages would make the *Pc* system more accessible and more importantly, largely independent of primates. In conclusion, this iPS-based *in vitro* system provides a promising alternative to investigate the dormant stage of malaria in the liver and may facilitate drug screening for compounds with activity against hypnozoites.

## Material and Methods

### Primary cells

Cynomolgus monkey fibroblasts used for iPS cell generation were obtained from ZenBio (monkey PDF091812) and Novartis (monkey 5501). Cynomolgus monkey primary hepatocytes were purchased from Biopredic. Human dermal fibroblasts used for iPS and hepatocyte generation were purchased from Invitrogen and human primary hepatocytes from KaLy-Cell. The donors provided written informed consents.

### Cynomolgus monkey iPS cell (cynIPS) generation

Primary cynomolgus monkey dermal fibroblasts were reprogrammed using Sendai viruses of CytoTune-iPS 2.0 Sendai Reprogramming Kit (Life Technologies, A16517) according to the standard protocol. Cells and viruses were plated on irradiated MEFs (Merck Millipore, PMEF-CF1) and cultivated with human embryonic stem cell medium (hESC) [KnockOut DMEM; 4.5 g/L glucose, 0.11 g/L sodium pyruvate, w/o glutamine (Gibco, 10829018), 20% KnockOut Serum Replacement (Gibco, 10828028), 1% glutaMAX (Gibco, 35050038), 1% MEM non-essential amino acids (NEAA) (Gibco, 11140035), 1% penicillin/streptomycin (Gibco, 15140-122), 10 ng/mL human basic fibroblast growth factor (bFGF) (Invitrogen, 13256029), and 0.1% β-mercaptoethanol (Gibco, 31350010)]. Colonies with hallmark of pluripotent morphology were readily visible between 2-3 weeks after transduction. These pluripotent colonies were picked and subcloned multiple times either on feeder cells or on Matrigel (BD Biosciences, 354277) coated plates in hESC or mTeSR1 medium (Stem Cell Technologies, 05850). This serial subcloning was performed until remnants of Sendai virus RNA were no longer detected and the morphology looked stable. As constant bFGF activity is essential for cynIPS maintenance, mTeSR1 medium was progressively replaced by StemFlex medium (Gibco, A3349401) which allows maintenance of bFGF levels overtime. Pluripotency was assessed by immunofluorescence staining, FACS analysis, and differentiation potential in all three germ layers (see sections below). Karyotype of each selected line was checked by Comparative Genomic Hybridization (aCGH method) (Cell Line Genetics, Madison, WI) and all showed a normal karyotype.

### Flow cytometry analysis for bFGF experiment and iPS characterization

CynIPS cells were dissociated with TryplE (Invitrogen, 12604), followed by fixation and permeabilization using Cytofix/Cytoperm reagent (BD Biosciences, 554655 and 51-20S1KZ) according to the manufacturer. Cells were stained with Nanog-PE (BD Biosciences, 560589a), Oct3/4-PerCP (BD Biosciences, 560794) and Sox2-A647 (BD Biosciences, 562139) intracellular primary antibodies for 45 min at 4 °C, washed twice with ice cold Cytoperm reagent and re-suspended in 1% fetal calf serum (FCS) in PBS (Invitrogen, 14190). Cells were subsequently analyzed on a CYAN flow cytometer (Beckmann Coulter).

### Karyotyping

For each tested cell line, 1.5E05 cells were seeded onto a T25 Matrigel coated flask in StemFlex medium supplemented with 10 µM rock inhibitor (Merck Millipore, Y-27632). Medium was refreshed the following day without rock inhibitor and lived cells were sent over to Cell Line Genetics when 20-30% confluence was reached. Karyotype of each selected line was checked by Comparative Genomic Hybridization (aCGH method).

### Differentiation potential

#### 1. Endoderm differentiation

Endodermal cells were induced from cynIPS cells using STEMdiff Definitive Endoderm (DE) Kit (Stem Cell Technologies, 05110). CynIPS cells were seeded onto a 24-well plate (TPP, 92024) coated with Matrigel and cultivated with StemFlex medium until culture reached 95-100% confluence. Cells were washed with PBS and medium was switched to STEMdiff medium with 1% DE supplements A and B. The next day, medium was switched to STEMdiff medium with 1% DE supplement B. Medium was changed every day. Cells were fixed after 5 days of differentiation with 4% paraformaldehyde (Electron Microscopy Sciences, 15714) in PBS and immunofluorescence staining was performed with FoxA2 (1:50 dilution, Abcam, ab60721) and Oct4 (1:1,000 dilution, Stemgent, 09-0023) primary antibodies.

#### 2. Ectoderm differentiation

Neuronal precursors were differentiated from cynIPS cells using a modified dual Smad inhibition protocol. 6E04 undifferentiated iPS cells were seeded onto a 96-well ULA (ultra low attachment) plate (Costar, 7007) in 0.1 mL StemFlex medium with 10 ng/mL penicillin/streptomycin and 10 μM rock inhibitor to prevent apoptosis. 24 hours after seeding, Embryoid Bodies (EBs) were formed and 0.1 mL fresh StemFlex medium were added to the wells. The next day, 30 EBs were transferred to a 4-well Matrigel coated plate with StemFlex medium (7 or 8 EBs seeded per well). 6 hours later, StemFlex medium was replaced by neural induction medium (20% KnockOut™ Serum Replacement, 0.1 mM MEM NEAA, 0.1 mM β-mercaptoethanol, 75% DMEM/F12/glutaMAX; 1% penicillin/streptomycin, 10 ng/mL human bFGF, 10 μM SB 431542 (Tocris, 1614), and 1 μM LDN 193189 (Stemgent, 04-0074). Medium was changed every other day until day 13. Cells were fixed after 14 days of differentiation with 4% paraformaldehyde in PBS for 10 min at room temperature (RT) and immunofluorescence staining was performed with βIII-tubulin (1:5,000 dilution, Sigma, T8660), NF200 (1:6,000 dilution, Abcam, ab72996), NeuN (1:500 dilution, Merck Millipore, ABN78), and Oct4 (1:1,000 dilution, Stemgent, 09-0023) primary antibodies.

#### 3. Mesoderm differentiation

6E04 undifferentiated cynIPS cells were seeded onto a 96-well ULA plate in 0.1 mL StemFlex medium with 10 ng/mL penicillin/streptomycin and 10 μM rock inhibitor to prevent apoptosis. 24 hours after seeding, EBs were formed and 0.1 mL fresh StemFlex medium were added to the wells. The next day, medium was replaced by STEMdiff Mesoderm Induction Medium (Stem Cell Technologies, 05220). Medium was refreshed daily until spontaneous EB contraction was observed after 10 days of differentiation (Movie 1).

### Generation of cynomolgus hepatocyte-like cells (cynHLCs)

CynIPS cells were harvested and seeded (1.1E05 cells/well) onto a 96-well plate (Greiner, 655090) coated with Laminin 521 (Biolamina, LN521) in StemFlex medium supplemented with 100 µg/mL geneticin (G418) (Gibco, 10131-035), and 10 µM rock inhibitor. The next day, cells were washed with PBS and differentiation was started (day 0) in Chemically Defined Medium (CDM) [50% IMDM (Invitrogen, 31980-030), 50% DMEM/F12 (HAM) (Invitrogen, A14625DJ), 1% Insulin-Transferrin-Selenium (ITS) (Invitrogen, 51500-056), 0.1% CD Lipid Concentrate (Invitrogen, 11905-031) and 2% BSA (25 %) (Sigma, A7979)] supplemented with 1 µg/mL doxycycline (Dox). The next day, medium was changed with CDM supplemented with 1 µg/mL Dox. On day 2, medium was switched to William’s E medium (no phenol red, Gibco, A1217601) with Primary Hepatocyte Maintenance Supplements (Gibco, CM4000) and 1 µg/mL Dox. Medium was refreshed the next day. On day 4, Dox induction was stopped and William’s E medium containing Primary Hepatocyte Maintenance Supplements was supplemented with 5 µM of a combination of compounds (Doramapimod, and inhibitors of DHODH and Raf) to maintain cynHLCs until 17 days of differentiation. Medium was refreshed daily.

### Generation of human hepatocyte-like cells (hHLCs)

Human iPS cells were harvested and seeded onto a 96-well Laminin 521 coated plate (8E04 cells/well) in mTeSR1 medium supplemented with 100 µg/mL G418 and 10 µM rock inhibitor. The next day (day 0), cells were washed with PBS and differentiated for 3 days in CDM supplemented with 1 µg/mL Dox (medium was refreshed daily). On day 3, medium was switched to William’s E medium with Primary Hepatocyte Maintenance Supplements and 1 µg/mL Dox. Medium was refreshed on day 4 and 5. On day 6, Dox induction was stopped and cells were further differentiated in William’s E medium containing Primary Hepatocyte Maintenance Supplements until day 25. Medium was refreshed every other day.

### Immunofluorescence analysis

Cells were fixed with 4% paraformaldehyde in PBS, permeabilized with Triton X-100 (Sigma, T8787) in PBS and stained with primary antibodies visualized with appropriate fluorescently labeled secondary antibodies (Alexa Fluor). Cultures were counterstained with the nuclear marker Hoechst (Invitrogen, 33342) and immunostained samples were imaged using a Zeiss LSM 700 microscope. Primary antibodies references can be found in Table S1.

### Periodic Acid-Schiff (PAS) staining (Sigma, 395-1KT)

Fixed samples were rinsed with deionized water and then placed in 0.5% periodic acid aqueous solution for 5 min at RT. Samples were then rinsed with running tap water for 3 min and quickly rinsed with deionized water before being placed in Schiff reagent solution for 15 min. The same rinsing steps as described above were used and samples were then counterstained with Hematoxylin for 2 min. Samples were rinsed with deionized water for 3 min. Samples were rinsed twice with 70% ethanol, twice with 96% ethanol and twice with 100% ethanol for 1 min each. Samples were covered with PBS before imaging.

### Oil Red O staining

Fixed samples were washed twice with deionized water and placed in 60% isopropanol for 5 min. 30 mL Oil Red O Stock Solution (60 mg Oil Red O (BioVision, K580-24-3) in 20 mL 100% isopropanol) was mixed with 20 mL deionized water and filtered using a 0.2 µm syringe filter to make Oil Red O Working Solution. Samples were incubated with working solution for 15 min and then washed two to five times with deionized water until excess stain was no longer apparent. Samples were counterstained with Hematoxylin (BioVision, K580-24-2) for 1 min. Samples were washed two to five times with deionized water before imaging.

### ELISA assays

To assess albumin secretion capacity of hepatocytes, cynIPS cells were harvested and seeded onto a 24-well Laminin 521 coated plate (7.5E05 cells/well) and differentiated as previously described. Culture medium was collected every 24 hours before the next medium change. ELISA assay was performed with the collected supernatants according to the provider standard protocol (Immunology Consultants Laboratory, E-80AL).

### Hepatocyte survival screen

CynIPS cells were harvested and 2.5E04 cells/well were seeded using a multidrop dispenser onto 384-well Laminin 521 coated plates (Corning, 3712) in StemFlex medium supplemented with 100 µg/mL G418, and 10 µM rock inhibitor. Cells were differentiated as described above until day 4. At day 4, Dox induction was stopped, medium was replaced by William’s E medium with Primary Hepatocyte Maintenance Supplements and cells were treated with the Novartis compound library in the Echo^®^ Liquid Handling platform (Labcyte Inc.). Cells were returned to overnight incubation at 37 °C, 5% CO_2_. Cells were treated with the Novartis compound library every other day after medium change. At day 13, bright field images of all plates were acquired with a high-throughput high-content imaging system (Operetta, Perkin-Elmer) and CellTiter-Glo luminescent cell viability assay (Promega, G7570) was performed. Luminescent readings were obtained using an EnVision multilabel plate reader (Perkin-Elmer) after 10 min incubation at RT. Generated data were analyzed in TIBCO Spotfire^®^ (TIBCO Software Inc.).

### RNA sequencing

Total RNA was isolated using the Direct-zol RNA MiniPrep Kit (Zymo Research, R2071) including on-column DNase digestion according to the manufacturer’s instructions. RNA sequencing libraries were prepared using the Illumina TruSeq RNA Sample Prep kit v2 (Illumina, RS-122-2001) and sequenced using the Illumina HiSeq2500 platform. Samples were sequenced to a length of 2 × 76 base pairs. Read pairs from cynomolgus iPS, HLCs, and primary hepatocytes were mapped to the *Macaca fascicularis* genome and the cynomolgus gene transcripts from Refseq by using an in-house gene quantification pipeline (Schuierer & Roma, 2016). The human genome (hg38) was used for the mapping of human HLCs, primary hepatocytes, and liver related cancer cell lines. Genome and transcript alignments were used to calculate gene counts based on Ensembl gene IDs.

#### Generation of *Plasmodium cynomolgi (Pc)* sporozoites

The Biomedical Primate Research Centre (BPRC) is an AAALAC-certified institute. All rhesus macaques (*Macaca mulatta*) used in this study were captive bred for research purposes and were housed at the BPRC facilities in compliance with the Dutch law on animal experiments, European directive 2010/63/EU, and with the Standard for Human Care and Use of Laboratory Animals by Foreign institutions, identification number A5539-01, provided by the National Institutes of Health (NIH). Prior to the start of experiments, all protocols were approved by the local independent ethical committee, according to Dutch law. For each batch of infected mosquitoes, one rhesus macaque was infected with 1E06 *Plasmodium cynomolgi* M strain blood stage parasites and bled at peak parasitaemia, after which the monkey was cured from the malaria infection by *IM* injection of 7.5 mg/kg chloroquine on three consecutive days. Approximately 1,200 female *Anopheles stephensi* mosquitoes strain Sind-Kasur Nijmegen (Nijmegen University Medical Centre St. Radboud, Department of Medical Microbiology) were fed with this blood using an *ex vivo* glass feeder system.

### Sporozoite infection of cynHLCs

Sporozoite inoculation of cynHLCs was performed at Novartis Institutes for Biomedical Research (NIBR, CH) according to the methods of Dembélé et al. (*5*). *Pc* sporozoites were kindly provided by Anne-Marie Zeeman (BPRC, NL) and shipped in Leibovitz L15 medium (Invitrogen, 11415-056) with 3% FCS and 2% penicillin/streptomycin at 4 °C to ensure good sporozoite infectiousness. Upon arrival, 4-days differentiated cynHLCs were infected with different sporozoite densities in 96-well plates. Cultures were kept at 37 °C in 5% CO_2_ with daily medium changes. To evaluate the development of *Pc* liver stages, cultures were fixed with 4% paraformaldehyde at the indicated time points.

### Immunofluorescence staining of LS parasites

Infected hepatocytes were fixed with 4% paraformaldehyde for 30 min, followed by overnight incubation at 4 °C with rabbit primary anti-*Pc*Hsp70 antibodies and rat primary anti-*Pc*UIS4 antibodies both diluted in 1% BSA and 0.3% Triton X-100 in PBS solution. Primary antibodies were kindly provided by Anne-Marie Zeeman (BPRC, NL). Subsequently, donkey secondary IgG Alexa Fluor^®^ 555-conjugated anti-rabbit antibodies (1:1,000 dilution, Invitrogen, A31572), chicken secondary IgG Alexa Fluor^®^ 594-conjugated anti-rat antibodies (1:1,000 dilution, Invitrogen, A21471) were added for three hours at RT. Nuclei were counterstained with Hoechst for 10 min at RT, samples were finally covered with PBS and viewed under an inverted microscope (Leica DMI6000 or Zeiss LSM 700). Image acquisition and post processing were performed with proprietary Leica or Zeiss software, LAS X or Zen 2012 respectively. Primary and secondary antibodies references can be found in Table 1.

### Statistical analysis

Data are presented as mean ± standard deviation (SD). “n” refers to biological replicates. P < 0.05 was considered statistically significant (ns, not significant (P > 0.05); *P < 0.05; **P < 0.01; ***P < 0.001; ****P < 0.0001). GraphPad Prism 7 (GraphPad Software) was used for statistical analyses. In Figure 3C, one-way ANOVA was used to compare the groups followed by Dunnett post hoc test.

## Acknowledgments

We thank Isabelle Fruh, Bettina Leonhard, Carole Manneville, and Annick Werner for continuous support with iPS technology; Nicole van der Werff, Ivonne Nieuwenhuis, and Lars Vermaat for mosquito dissection and sporozoite preparation; Annemarie Voorberg-van der Wel for helpful discussions regarding infection assay and critical reading of the manuscript; Isabelle Claerr for technical assistance with high content imaging; Olaf Galuba for support in conducting the survival screen; Dominic Trojer for help in microscopy; Volker Heussler for insightful discussions; Benoit Fischer and Marianne Uteng for providing RNA samples from cynomolgus primary hepatocytes; Judith Knehr and Walter Carbone for RNA-sequencing; and the Walter Fischli-Foundation, the Bill and Melinda Gates Foundation and the Medicines for Malaria Venture for financial support.

## Author contributions

M.M., M.R., and P.M.: Conception and design, data analysis and interpretation, manuscript writing; M.P.: Conception and design, collection and assembly of data, data analysis and interpretation, manuscript writing; A.M.Z.: Provision of sporozoites, data analysis and interpretation; G.R., S.S.: Analysis of RNA-seq experiment; A.M.Z., C.H.M.K., E.L.F., G.R., L.K.A., M.M., M.R., P.M., S.A.M., S.S., and T.D.: Final approval of manuscript.

## Data and materials availability

All data are available in the main text or the supplementary materials.

