## Supplementary Information for "Persistence of *Plasmodium cynomolgi* hypnozoites in cynomolgus monkey iPS-derived hepatocytes"

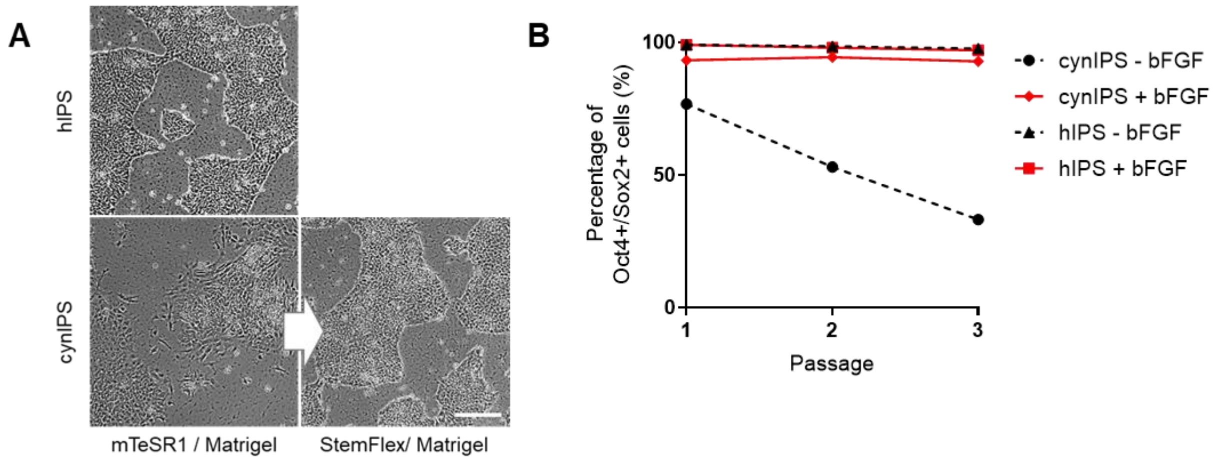


**Figure 1-figure supplement 1: Cynomolgus vs. human iPS cell maintenance after generation.** **A,** Brightfield images of human (h) and cynomolgus (cyn) iPS cells. CynIPS spontaneously differentiated when cultivated with mTeSR1 medium. Switching to StemFlex medium with stabilized bFGF was essential for cynIPS cell maintenance. Scale bars, 200 µm. **B,** FACS analysis of human and cynomolgus iPS cells (cultivation +/- bFGF) with Oct4 and Sox2 pluripotent markers.

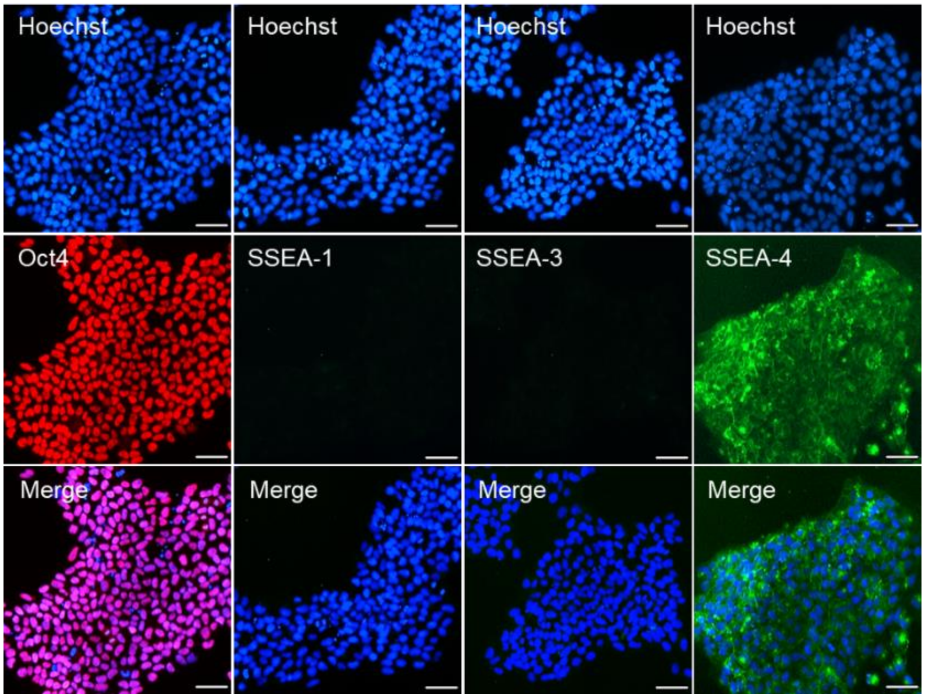


**Figure 1-figure supplement 2: Selected clone #13 from monkey PDF091812.** Immunofluorescence analysis of Oct4 (red), SSEA-1, SSEA-3, and SSEA-4 (green) in cynIPS cells. Nuclei were counterstained with Hoechst (blue). Scale bars, 50 µm.

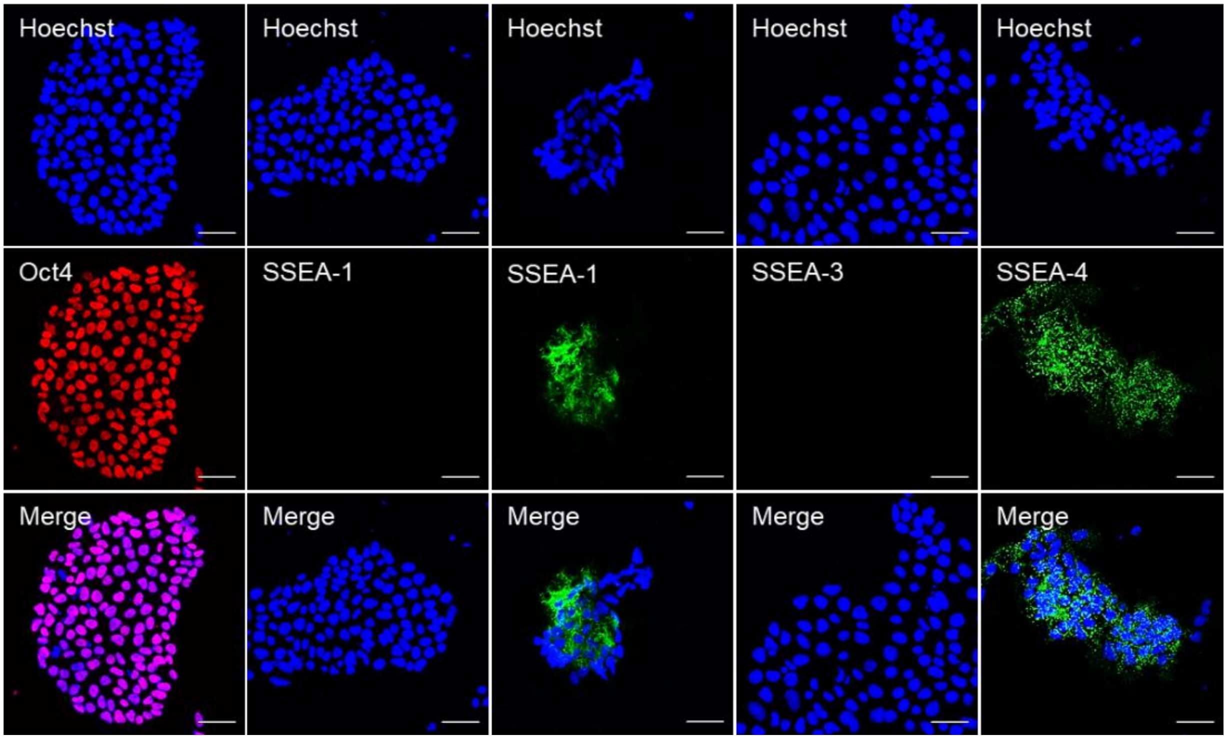


**Figure 1-figure supplement 3: Example of a clone not selected based on SSEA-1 expression.** Immunofluorescence analysis of an iPS clone from monkey 5501 of Oct4 (red), SSEA-1, SSEA-3, and SSEA-4 (green). Nuclei were counterstained with Hoechst (blue). Scale bars, 50 µm.

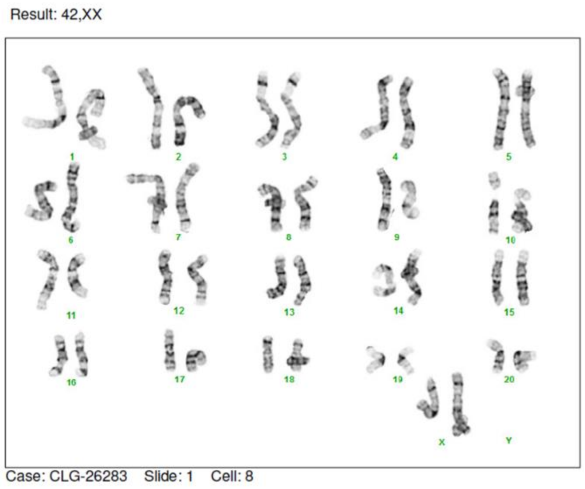


**Figure 1-figure supplement 4:** Karyotype analysis of cynIPS cells from clone #13 by Comparative Genomic Hybridization (aCGH method).

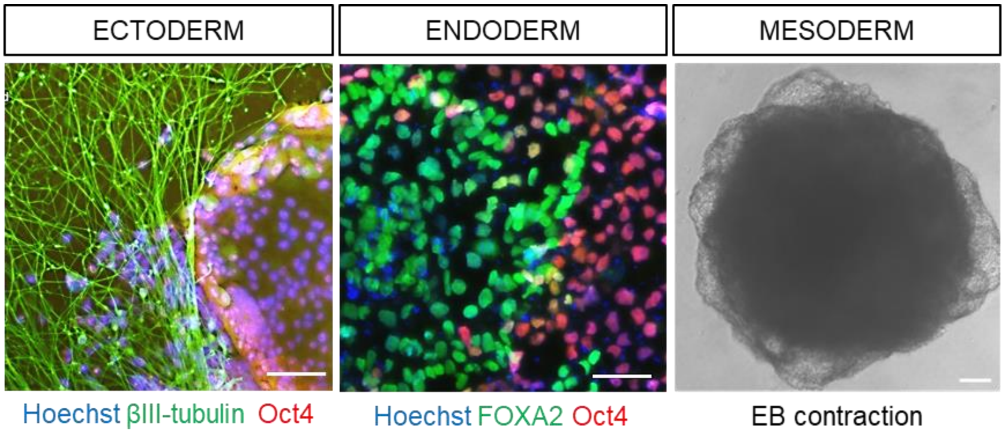


**Figure 1-figure supplement 5: Differentiation potential of cynIPS cells from clone #13.** Upon stimulation, cells expressed βIII-tubulin (green, ectoderm marker), FOXA2 (green, endoderm marker) and contracted spontaneously after embryoid body (EB) formation (mesoderm). Non-responsive cells remained pluripotent and expressed Oct4 (red). Nuclei were counterstained with Hoechst (blue). Scale bars, 100 µm.

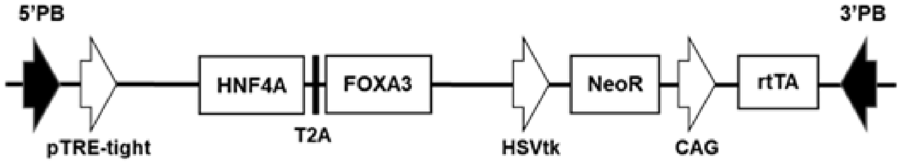


**Figure 1-figure supplement 6: Schematic representation of the Dox-inducible construct.** PB: PiggyBac DNA transposon; TRE: TET responsive element; HNF4A: Coding sequence of hepatocyte nuclear factor 4 alpha; FOXA3: Coding sequence forkhead box protein A3); T2A: self-cleavage peptides from *Thosea asigna* virus; HSVtk: Minimal promoter fragment from the HSV thymidine kinase (TK) promoter; NeoR: neomycin resistance gene; CAG: chicken β-actin promoter; rtTA: TET transactivator protein gene.

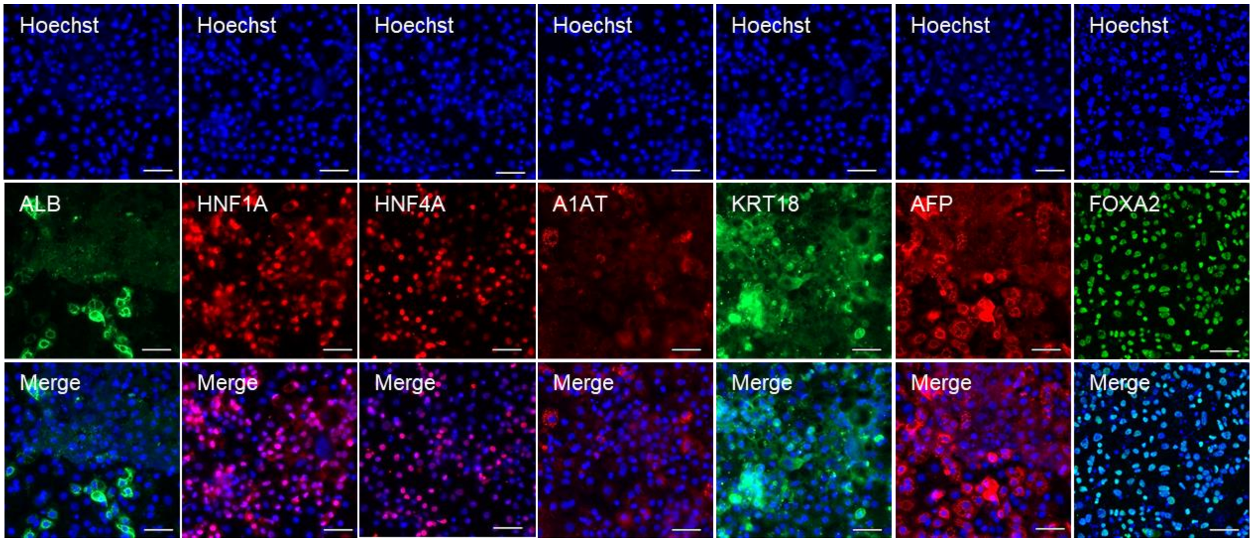


**Figure 1-figure supplement 7: Hepatic differentiation of clone #13.6 showed less Albumin staining.** Immunofluorescence analysis of cynHLCs after 6 days of differentiation with antibodies to HNF1A, HNF4A, AFP, A1AT (red), ALB, KRT18, and FOXA2 (green). Nuclei were visualized with Hoechst (blue). Scale bars, 50 μm.

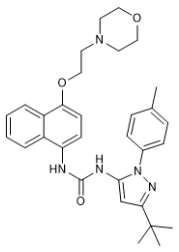

Doramapimod

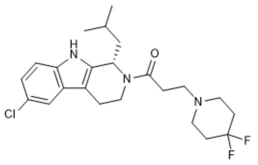

DHODH inhibitor

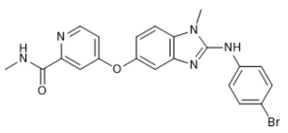

Raf inhibitor

| Compound name | IUPAC name | Canonical Smiles |
|---|---|---|
| Doramapimod | 1-[5-tert-butyl-2-(4-methylphenyl)pyrazol-3-yl]-3-[4-(2-morpholin-4-ylethoxy)naphthalen-1-yl]urea | CC1=CC=C(C=C1)N2C(=CC(=N2)C(C)(C)C)NC(=O)NC3=CC=C(C4=CC=CC=C43)OCCN5CCOCC5 |
| DHODH inhibitor | 1-[(1S)-6-chloro-1-(2-methylpropyl)-1,3,4,9-tetrahydropyrido[3,4-b]indol-2-yl]-3-(4,4-difluoropiperidin-1-yl)propan-1-one | CC(C)CC1C2=C(CCN1C(=O)CCN3CCC(CC3)(F)F)C4=C(N2)C=CC(=C4)Cl |
| Raf inhibitor | 4-[2-(4-bromoanilino)-1-methylbenzimidazol-5-yl]oxy-N-methylpyridine-2-carboxamide | CNC(=O)C1=NC=CC(=C1)OC2=CC3=C(C=C2)N(C(=N3)NC4=CC=C(C=C4)Br)C |

**Figure 3-figure supplement 1:** Chemical structures of the hit compounds identified in survival screen (Doramapimod, DHODH, and Raf inhibitors) and table displaying the IUPAC names and the canonical smiles of each compound.

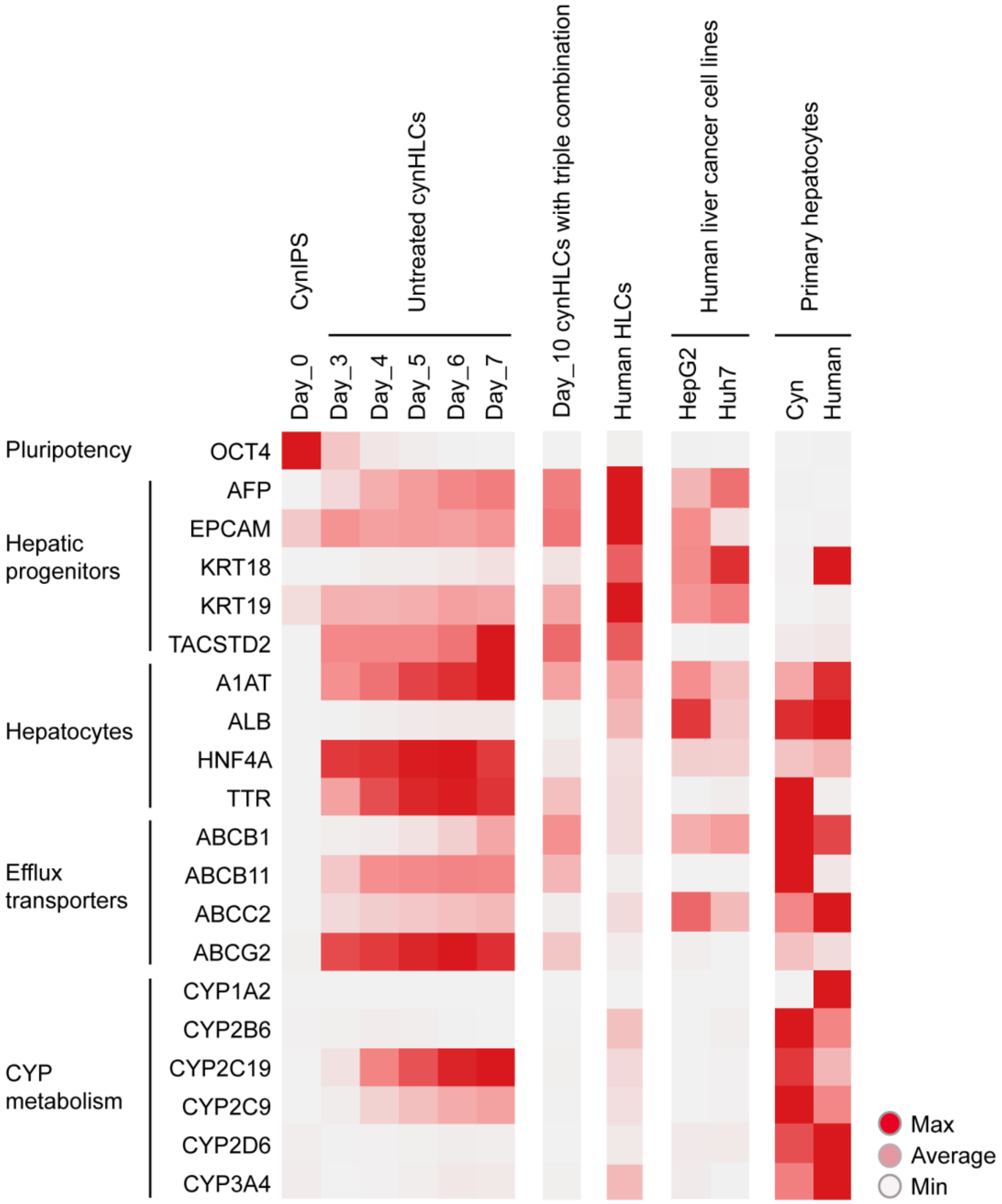


**Figure 3-figure supplement 2:** Heatmap of the RNA-seq based mean expression of cynIPS cells (Day_0), cynHLCs (Day_7 untreated and Day_10 triple combination), and human HLCs for selected genes indicative of cell types (Pluripotency, Hepatic progenitors, and Hepatocytes) or cellular processes (Efflux transporters and CYP metabolism). Expression values in transcripts per million (tpm) are colored relative to the maximum (red) and minimum (white) expression value for each gene.

| Name | Symbol | Provider | Reference | Specie | Dilution |
|---|---|---|---|---|---|
| Alpha-1-antitrypsin | A1AT | Dako | A0012 | Rabbit | 1:10,000 |
| Alpha-1-Fetoprotein | AFP | Dako | A0008 | Rabbit | 1:500 |
| Albumin | ALB | Bethyl | A80-129A | Goat | 1:500 |
| Class III β-tubulin | βIII-tubulin | Sigma | T8660 | Mouse | 1:5'000 |
| Cluster of Differentiation 81 | CD81 | Invitrogen | MA5-13548 | Mouse | 1:100 |
| Forkhead box protein A2 | FOXA2 | Abcam | ab60721 | Mouse | 1:50 |
| Cytokeratin 18 | KRT18 | Abcam | ab82254 | Mouse | 1:100 |
| Hepatocyte nuclear factor 1A | HNF1A | Cell Signaling Technologies | 12425 | Rabbit | 1:100 |
| Hepatocyte nuclear factor 4A | HNF4A | Cell Signaling Technologies | 3113 | Rabbit | 1:100 |
| Octamer-4 | Oct4 | Stemgent | 09-0023 | Rabbit | 1:1,000 |
| *P. cynomolgi* 70 kilodalton heat shock proteins | Pc-Hsp70 | Anne-Marie Zeeman, BPRC | 20285 | Rabbit | 1:1,000 |
| *P. cynomolgi* up-regulated in infective sporozoites gene 4 | Pc-UIS4 | Anne-Marie Zeeman, BPRC | 5262527 | Rat | 1:400 |
| Scavenger Receptor class B member 1 | SRB1 | Novus Biologicals | NB400-104 | Rabbit | 1:100 |
| Stage-specific embryonic antigen-1 | SSEA-1 | Chemicon/Milipore | MAB4301 | Mouse | 1:200 |
| Stage-specific embryonic antigen-3 | SSEA-3 | Stemgent | 09-0014 | Rat | 1:200 |
| Stage-specific embryonic antigen-4 | SSEA-4 | Stemgent | 09-0006 | Mouse | 1:1,000 |
| Alexa Fluor 555 donkey anti-rabbit | Dk@Rb | Invitrogen | A31572 | Donkey | 1:1,000 |
| Alexa Fluor 488 donkey anti-goat | Dk@Gt | Invitrogen | A11055 | Donkey | 1:1,000 |
| Alexa Fluor 488 goat anti-mouse | Gt@Ms | Invitrogen | A10667 | Goat | 1:1,000 |
| Alexa Fluor 594 chicken anti-rat | Ch@Rt | Invitrogen | A21471 | Chicken | 1:1,000 |
| Goat anti-rabbit IgG-FITC | Gt@Rb | Anne-Marie Zeeman, BPRC | | Goat | 1:200 |
| Hoechst | Hoechst | Invitrogen | 33342 | N.A. | 1:10,000 |

**Table 1:** Primary and secondary antibodies used for immunofluorescence analysis.

**Movie 1:** Embryoid body (EB) contraction occurred spontaneously after 10 days of differentiation in ultra low attachment plate.
